# FiPhoPHA - A fiber photometry python package for post-hoc analysis

**DOI:** 10.1101/2025.06.12.659422

**Authors:** Vasilios Drakopoulos, Alex Reichenbach, Romana Stark, Claire J. Foldi, Philip Jean-Richard-dit-Bressel, Zane B. Andrews

## Abstract

Fiber photometry is a neuroscience technique that can continuously monitor *in vivo* fluorescence to assess population neural activity or neuropeptide/transmitter release in freely behaving animals. Despite the widespread adoption of this technique, methods to statistically analyse data in an unbiased, objective, and easily adopted manner are lacking. Various pipelines for data analysis exist, but they are often system-specific, only for pre-processing data, and/or lack usability. Current post hoc statistical approaches involve inadvertently biased user-defined time-binned averages or area under the curve analysis. To date, no post-hoc user-friendly tool with few assumptions for a standardised unbiased analysis exists, yet such a tool would improve reproducibility and statistical reliability for all users. Hence, we have developed a user-friendly post hoc statistical analysis package in Python that is easily downloaded and applied to data from any fiber photometry system. This **Fi**ber **Pho**tometry **P**ost **H**oc **A**nalysis (**FiPhoPHA**) package incorporates a variety of tools, a downsampler, bootstrapped confidence intervals (CIs) for analyzing peri-event signals between groups and compared to baseline, and permutation tests for comparing peri-event signals across comparison periods. We also include the ability to quickly and efficiently sort the data into mean time bins, if desired. This provides an open-source, user-friendly python package for unbiased and standardised post-hoc statistical analysis to improve reproducibility using data from any fiber photometry system.

## Introduction

Fiber photometry is a tool used by neuroscientists to measure population level neural activity or region- and/or cell-specific neuropeptide/transmitter release in freely moving animals (Kielbinski and Bernacka, 2024). The rapid rise in popularity of this technique is driven by the ever-expanding toolbox of genetically engineered proteins that emit fluorescent light in response to changes in intracellular calcium (a proxy for neuronal activity) or neuropeptide and neurotransmitter dynamics (Simpson et al., 2024). The most well-known of these is the genetically-encoded green calcium indicator, GCaMP, which is used to track neuronal activity based on changes in intracellular calcium (Lütcke et al., 2010). Typically, blue light (∼465nm; experimental) and violet light (∼405nm; isosbestic control) is sent into the brain through a fiber optic cable attached to a fiber optic implant fixed in the brain. In this way, wavelength-specific excitation of cells expressing GCaMP (or other sensors excited at 465nm) emit light, which are then passed through a filter and detected on a photodetector. Other sensors fluoresce at other wavelengths, such as red-shifted calcium indicators, such as jRGECO and sRGECO and dopamine (GRAB-rDA) sensors that have excitation wavelengths of ∼560nm (Dana et al., 2016; Fenno et al., 2020; Zheng et al., 2024).

Before statistical comparisons can be made, several data pre-processing steps are required to artifact-correct the neural dynamic-tracking experimental signal. Typically, the isosbestic control signal is fit to the experimental signal, as applied in the equation 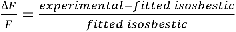 to account for neural dynamic-independent changes in fluorescence (e.g., motion artifacts, photobleaching). Following this, a common approach is to normalize the signal via z-scoring, where 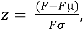 with Fµ and Fσ being the mean and standard deviation of a reference period. Various pipelines for pre-processing data from various fiber photometry systems have been made freely available for use (Akam and Walton, 2019; Bruno et al., 2021; Sherathiya et al., 2021; Conlisk et al., 2023; Murphy et al., 2023; Bridge et al., 2024).

From here, post hoc statistical comparisons can be made of peri-event signals using a variety of approaches. Peri-event analyses is where data is aligned to an experimental event, to assess how the data changes relative to this event. This includes user-defined summary statistics for peri-event time bins, such as area under the curve, average z-score, or peak z-score (Lerner et al., 2015; de Jong et al., 2019; Clarke et al., 2023; de Araujo Salgado et al., 2023; So et al., 2023; Reichenbach et al., 2024). However, these approaches are inadvertently biased based on user-defined time periods for analysis with potential unmet assumptions about data distribution. Thus, an easy-to-use post hoc analysis tool with fewer statistical assumptions independent from any temporal information and greater power is desperately needed to improve reliability and data reproducibility in an unbiased manner.

To address this, a previous study (Jean-Richard-dit-Bressel et al., 2020) compared alternative analysis approaches including parametric confidence intervals (CIs), non-parametric bootstrapped CIs, and permutation tests that did not require obtaining summary statistics from arbitrary time bins. These approaches permit data-driven, temporally precise identification of when peri-event signals differ from baseline or comparison signals. Furthermore, bootstrapped CIs and permutation tests are forms of resampling methods that do not require certain assumptions about the population distribution to be met (e.g., that signals are normally distributed). A key finding of this study was that Type 1 error (false positives) could be adequately reduced without inflating Type 2 error (false negatives) by using a consecutive threshold for significance, with this threshold being critical for the validity of these techniques.

Although this study concluded bootstrapped CIs and permutation tests have highly desirable qualities for analyzing peri-event waveforms, there are no user-friendly, computationally simple, freely available packages for deploying these methods in open source programming language like Python (existing packages [https://github.com/philjrdb/FibPhotom] have only been made available for MATLAB, a proprietary program). Recently, another pipeline for post-hoc analysis was published, however this analysis relies on data distribution assumptions that may not accurately reflect the non-parametric, noisy, and non-linear dynamics of a fiber photometry signal (Loewinger et al., 2025). Moreover, this approach is not user-friendly and requires data modelling, which can be computationally demanding across larger datasets. Therefore, we developed a downloadable Python package, FiPhoPHA (<u>Fi</u>ber <u>Pho</u>tometry <u>P</u>ost <u>H</u>oc <u>A</u>nalysis) that does not require a strong coding background to implement. FiPhoPHA serves as post hoc photometry analysis toolkit for all photometry users to improve reliability and data reproducibility in an unbiased manner. This package includes a downsampler, popular time bin analyses, bootstrapped CIs, and permutation tests.

## Methods

In the following methods and instructions section, we frequently refer to terms such as “number of resamples”, “number of permutations” and “consecutive threshold”. We define these terms here for clarity.

Number of Resamples: The number of times random samples will be drawn with replacement to determine bootstrapped confidence intervals.

Number of Permutations: The amount of random shuffling of data to determine p-values for permutation tests.

Consecutive Threshold: The number of individual data points in the dataset that need to be in a row to consider the data statistically signficant.

The package is published and available at PyPI: https://pypi.org/project/fiphopha/. Individual files are also available on github: https://github.com/VasiliDrakopoulos/fiphopha.

1. **Create a virtual environment in any terminal of Python (or if you have Anaconda installed):**

python -m venv ( environment name) or conda create –n ( environment name)

2. **Install package:**

pip install fiphopha

3. **Run main script:**

fiphopha

The following text will appear in the terminal where the user can pick 1-4 for the appropriate analysis:

**Box 1.**
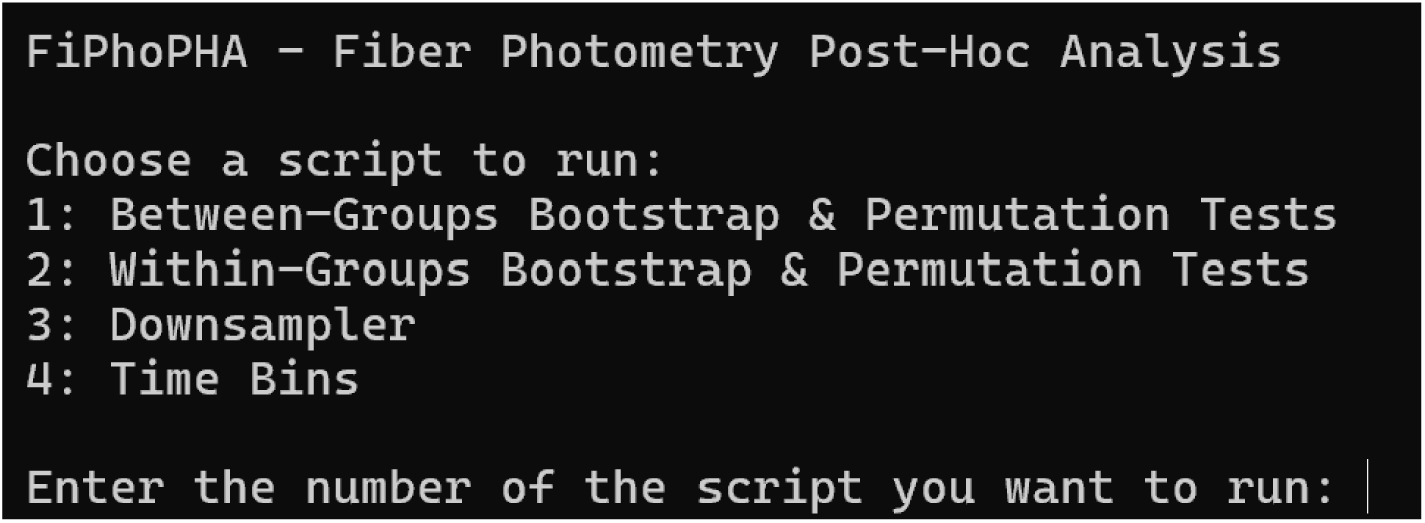

### Downsampler

Downsampling reduces the amount of data used in analysis. This is particularly useful when the original sampling rate is much greater than that needed to capture neural dynamics. Different researchers downsample their signal from pre-processing by different factors, this downsampler can be coupled with other post-hoc analysis to streamline processes, whilst still providing the temporal resolution wanted by the researcher. This downsampler works by averaging every n^th^ data point chosen. We recommend downsampling a fiber photometry signal according to the kinetics of the sensor and computational demand. If a signal is over downsampled, there is risk of losing temporal resolution and less data points for statistical comparison. But for recordings that last several minutes to hours this may be appropriate to offset computational demand.

### Mean time bins

Although previously described open-source fiber photometry analyses have the potential to conduct this analysis (Bridge et al., 2024), we have included this as an additional option to aid researchers in analysing their data. This calculates the mean across specified bins, which could include a baseline period and subsequent specific time points during the comparison period after an event.

### Bootstrapped confidence intervals and permutation tests

These analysis approaches were adapted from Jean-Richard-dit-Bressel and colleagues (Jean-Richard-dit-Bressel et al., 2020). They provide temporally specific alternatives to simple time bin analyses (e.g., mean time bins, Area Under the Curve), as it reports when peri-event dynamics significantly differ from the null hypothesis. Our toolkit is set up to run bootstrapped CIs at every timepoint in the peri-event window to determine statistical significance. CIs are a range estimate of the true population value (e.g., the true mean fluorescence change value at event onset). Bootstrapping obtains these CIs by repeatedly resampling the data with replacement and calculating a bootstrapped estimate each time. If performed a sufficient number of times (e.g., 1000 times), the resulting distribution of bootstrapped estimates, created from representative recombination of the dataset, serves as an approximation of the true sampling distribution and a range for our estimate can be obtained from percentiles of this bootstrap distribution (Efron and Tibshirani, 1994). It should be noted that increasing the number of bootstraps or permutations does not narrow the CI. Rather it provides a more representative bootstrapped distribution, which asymptotes quickly, hence 2000 bootstraps may not necessarily be better than 1000 (Jean-Richard-dit-Bressel et al., 2020). Given this bootstrapped CI does not factor sample size, an adjustment of 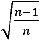 has been incorporated to account for small sample narrowness bias (Hesterberg, 2015). The output provides a binary significance map; 1 denotes significance and 0 denotes no significance for a given peri-event timepoint, for comparisons of each group to baseline this is determined by whether the CIs include the null of 0. There are also options for between or within-subjects analysis that includes different statistical analysis. For significance for between-subjects, the distributed means of each group are bootstrapped and subtracted. The low and high CIs from the bootstrapped distributed difference are calculated and determined if they include the null of 0. For within-subjects group comparison, each individual animal/trial is subtracted between groups and the product of the difference is bootstrapped. Due to this, for within-subjects, the same number of animals/trials in each group is needed. CIs are computed from the bootstrapped difference and difference is determined if they include the null of 0. The choice of analysing by animal or trial depends on the researcher’s specific question. We note that unbalanced trial data can inflate the reliability of an effect but may be appropriate in certain circumstances (Editorial, 2018). Hence it is important for researchers using this package to choose between or within-subjects comparisons using animal/trial data according to their specific data and the analysis required.

Permutation tests are another form of non-parametric resampling test that evaluates whether two groups are similar. It provides an exact p-value for the observed difference between groups to have come from the same distribution. This compares over a distribution of data between groups for a specified period. Hence, we have incorporated in the output file of FiPhoPHA, a sheet that provides p-values per timepoint between comparison periods. This non-parametric test makes few assumptions about the underlying distributions of comparison periods (Pesarin and Salmaso, 2010). These analyses have been used to compare neural dynamics signals between different events, and events to a reference baseline period (Pascoli et al., 2018; Ghareh et al., 2022). It is important to note that researchers often choose different baseline periods surrounding an event, usually a timeframe before the event or the entire recording period, which will likely change the y axis range considering that the neural data obtained during recording represents a relative change. This may change the results as depending on the relative change of the data, the signal may deviate differently from baseline.

### Choosing a consecutive threshold for significance

A consecutive threshold (also known as a temporal threshold) stipulates a minimum period that needs to be continuously significant for a comparison to be considered significant. This threshold is needed due to the issue of multiple comparisons and random signal crossings when analysing all timepoints in the peri-event window (e.g. through bootstrap CI or permutation tests). This consecutive threshold effectively combats Type 1 errors (false positives) without substantially increasing Type 2 errors (false negatives) (Jean-Richard-dit-Bressel et al., 2020). The appropriate consecutive threshold depends on several factors, including sensor kinetics and how a signal has been processed (e.g., low-pass filtering).

Sensor kinetics refers to how quickly a sensor’s signal can change in response to the dynamic it tracks (i.e., rise and decay time). Rise and decay times indicate the shortest duration signal that a sensor reports and therefore represents a logical temporal lower-bound for what is occurring physiologically. For example, in our demonstration of the FiPhoPHA package provided below, we used a dopamine sensor (GRAB-DA1m) that has an estimated on (rise) time of ∼0.08 seconds and an off (half-decay) time of ∼0.33 seconds (Sun et al., 2018). Therefore, real changes in dopamine concentrations are unlikely to translate to deviations in GRAB-DA signal that occur for less than 0.41 seconds. So, 0.41sec can serve as a kinetics-based consecutive threshold that allows identification of real signal change, while limiting the detection of spurious signal change (Type 1 errors). In our example data below, we have used 400ms for our consecutive threshold based on sensor kinetics, which is equal to 4 significant data points in a row when sampled at 10 Hz (referred to as consecutive threshold 4).

Low-pass filtering is used to improve signal-to-noise by removing the high-frequency signals, often associated with noise. Therefore, a consecutive threshold needs to account for any low pass filtering applied to photometry data. Previous studies observed that a consecutive threshold of 1sec/filter Hz (the “low-pass frequency window”) was an effective rule-of-thumb for reducing family-wise Type 1 error (Jean-Richard-dit-Bressel et al., 2020). In the FiPhoPHA demonstration below, our photometry system applied a 6 Hz low-pass filter during recordings, so a consecutive threshold based on the low-pass filter should be greater than 1/6sec (∼170 ms). To account for this in our example data below, we have used 200ms for our low-pass filter consecutive threshold, which is equal to 2 significant data points in a row when sampled at 10 Hz (referred to as consecutive threshold 2).

A more conservative consecutive threshold could also be chosen that considers other factors such as the time taken for user-defined behavioral classification (ie the time taken to perform or observe and classify a change in behavior). It is important to acknowledge that this used-defined threshold is harder to objectively define when compared to a consecutive threshold based on low-pass filtering or sensor kinetics and may not always be appropriate. In our example data below, we have user-defined consecutive threshold of 500 ms, which is equal to 5 significant data points in a row when sampled at 10 Hz (referred to as consecutive threshold 5). Our user-defined threshold of 500 ms is based on the time required for a mouse to collect and consume a 20 mg sucrose pellet.

Given multiple factors can be used to determine a consecutive threshold, we recommend choosing the more conservative (longer duration) threshold between the kinetics of the sensor and the low-pass filter frequency. For the examples described above, the temporal threshold based on kinetics (400 ms: consecutive threshold 4 when data are sampled at 10 Hz) should be chosen over the low-pass filter threshold (200 ms: consecutive threshold 2 when data are sampled at 10 Hz). To show how temporal thresholds influence results, we have presented data at 95% and 99% CI with and without the consecutive thresholds described above (Figure 2 and 3).

Importantly, the data sampling frequency should be considered when applying a consecutive threshold to downsampled data. For example, applying a consecutive threshold of 10 to data sampled at 10Hz (ie 1 data point every 100ms, consecutive threshold is 10 x 100ms), would be the same as a consecutive threshold of 1 when downsampled to 1Hz (ie 1 data point every 1000ms, 1 x 1000ms).

## Instructions

The following protocol is to install the FiPhoPHA python package and provides instructions on the setup by using example data from Reichenbach and colleagues, randomly selected data that uses three groups of data (Figure 2H) (Reichenbach et al., 2022). The example uses the within-subjects analysis as it uses the same animals for three trials/events of dopamine release in the nucleus accumbens from a non-food object, chow, and peanut butter. A windows, mac, or linux computer operating system is required with python installed https://www.python.org/downloads/ and Microsoft Excel https://www.microsoft.com/en-au/microsoft-365/excel.

For all analyses arrange data in excel as follows:

Each sheet in the excel document represents a group of data and each column is a different animal/trial as seen in the red and green in below. The timepoints of the dataset are now represented by the row index of the excel document.

**Box 2.**
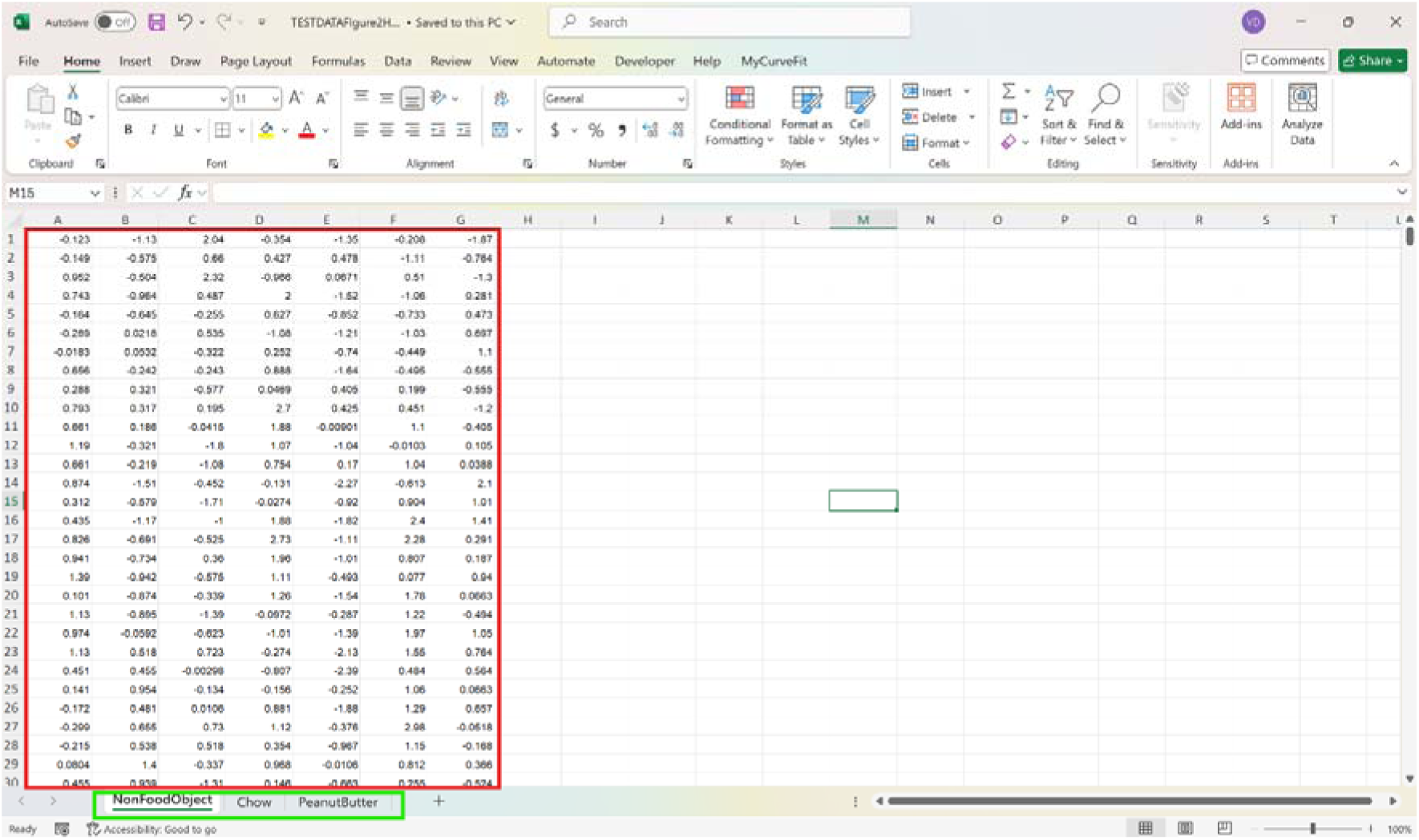

### Downsampler

1. Run fiphopha script and select 3: Downsampler.

2. The following graphical user interface (GUI) will be presented:

**Box 3.**
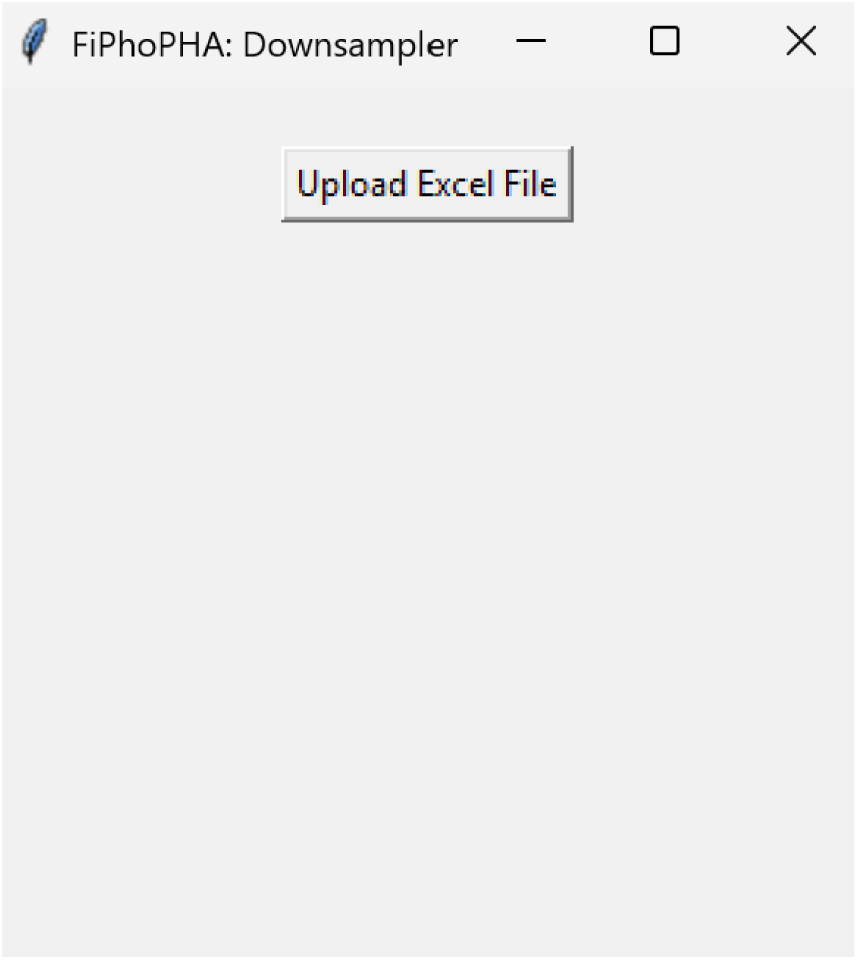

3. Press upload excel file and select your excel file.

4. The following GUI will be presented for the downsample factor to be selected:

**Box 4.**
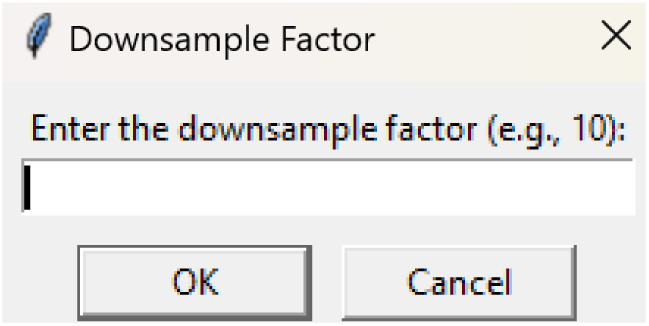

Since the following example data is sampled every 99.8ms (∼10 Hz), we will provide an example of downsampling by a factor of 10, to get an output of ∼1Hz.

5. The following output file saved as the original excel document name_downsampled_10 will show in the same directory as the input file.

**Box 5.**
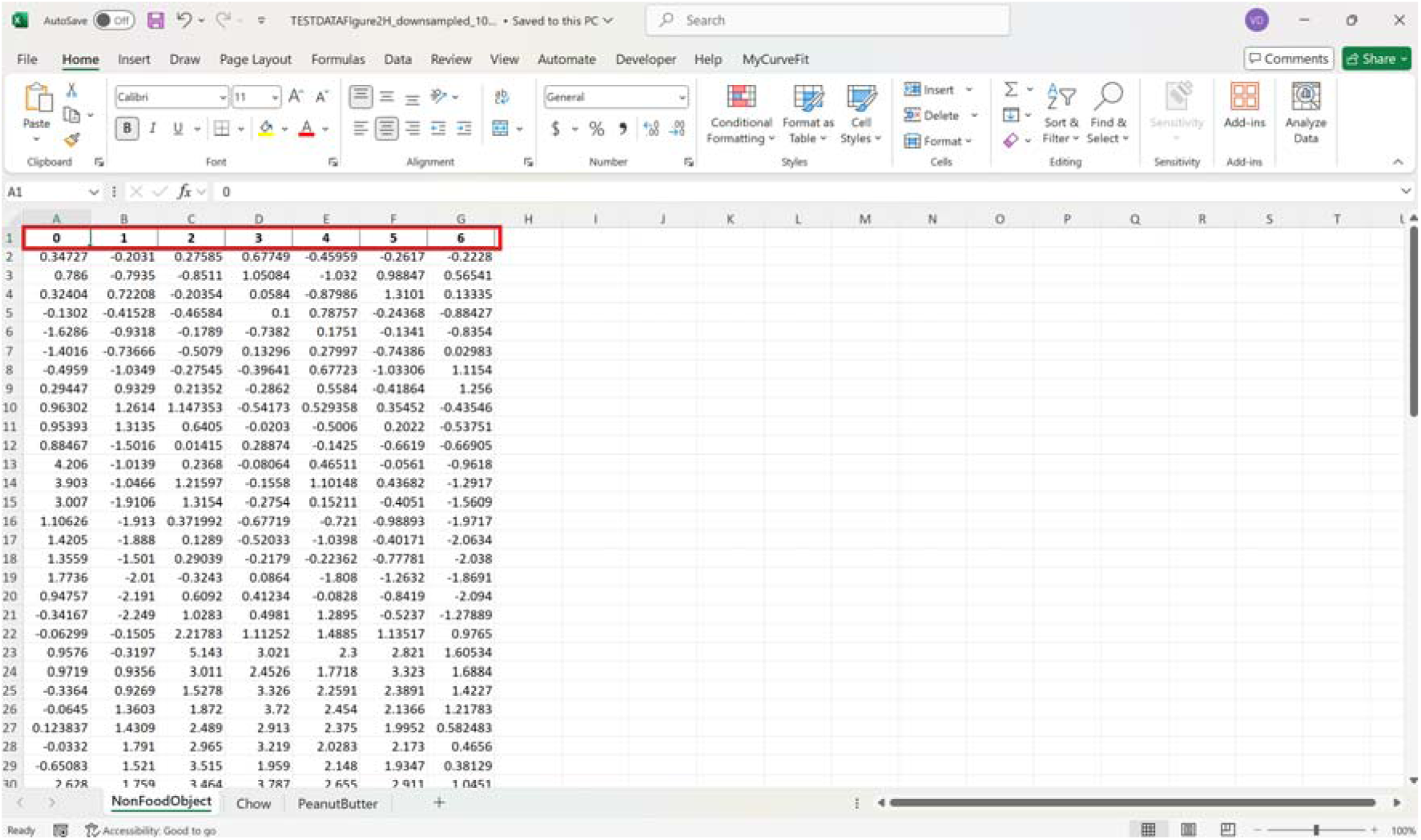

The red box shows the indexed columns of the output file to specify how many animals/trials the code analysed per sheet.

### Mean time bins

1. Run fiphopha script and select 4: Time bins.

2. The following GUI will be presented:

**Box 6.**
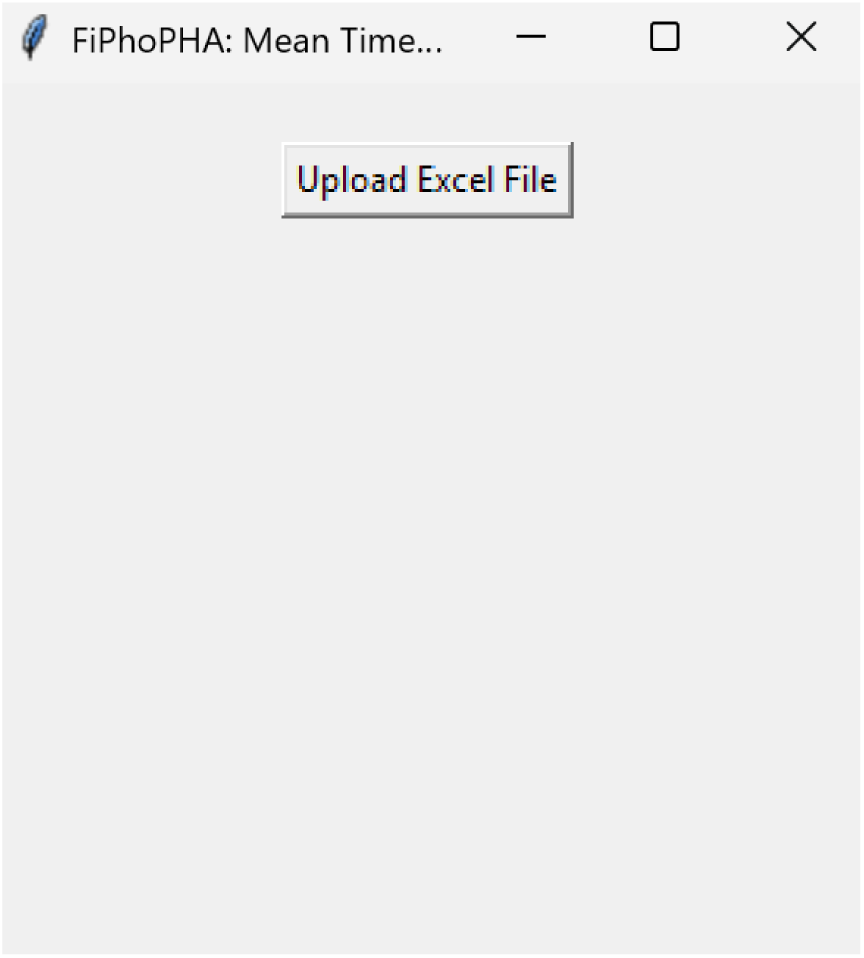

3. Press upload excel file and select your excel file.

4. The following GUI will be presented for the number of time bins. Since the example data used presents 813 time points between -20 seconds before event and 60 seconds after event, we have selected four time bins of approximately 20 seconds each. In this example, our data is sampled at 10 Hz so the time bins for 20 second intervals are 0-212, 213-416, 417-619, and 620-812 for the excel row index. The max value is 1 less than the max value due to python conventions of 0 being excel row index 1.

**Box 7.**
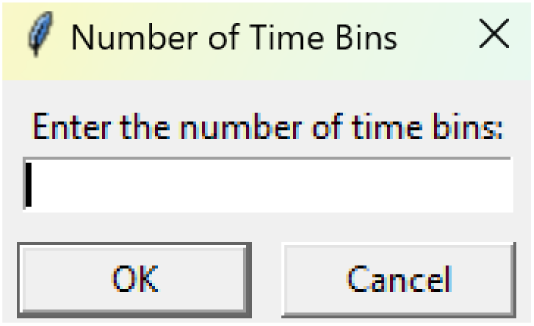

5. After number of time bins are selected, the following GUI is presented to enter the start row of the 1^st^ time bin.

**Box 8.**
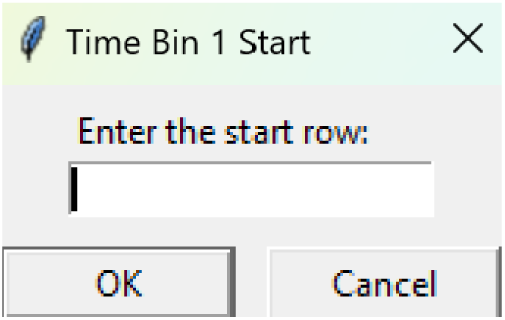

6. After, the following GUI is shown to enter the end row of the 1^st^ time bin.

**Box 9.**
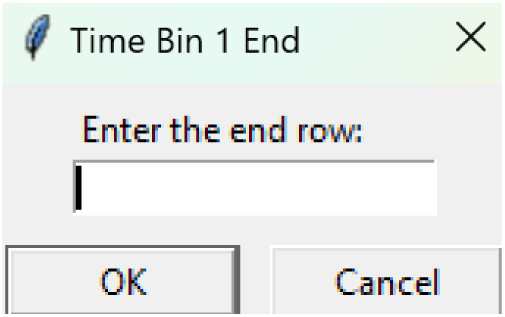

7. Repeat steps 5 to 6 for each time bin until all are entered

8. After all are entered, an output excel file will be saved to the same directory as the input file with the title of the input file with name_time_bin_means.

**Box 10.**
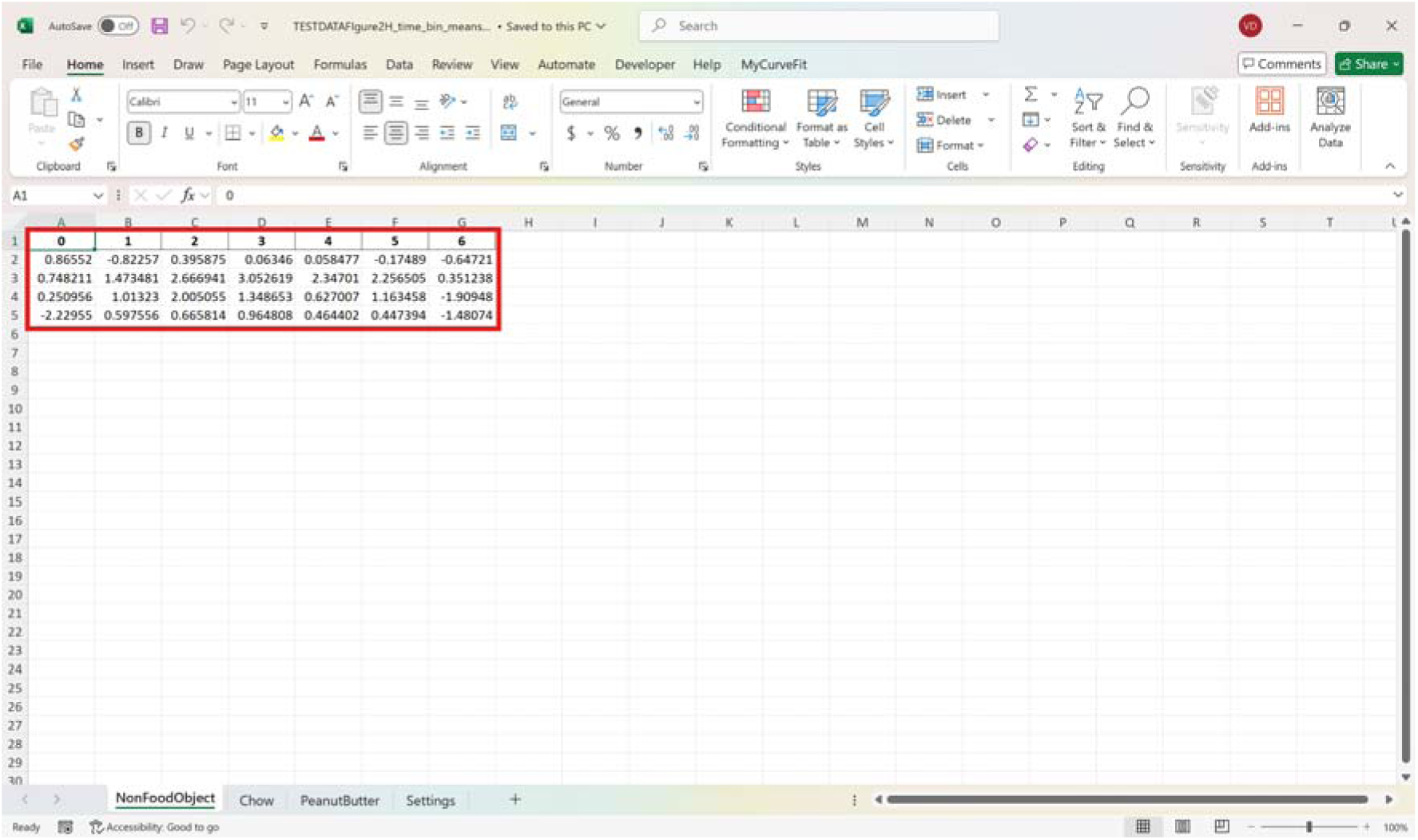

The red box shows the indexed columns of the output file to specify how many animals/trials the code analysed per sheet with the mean time bins underneath.

**Box 11.**
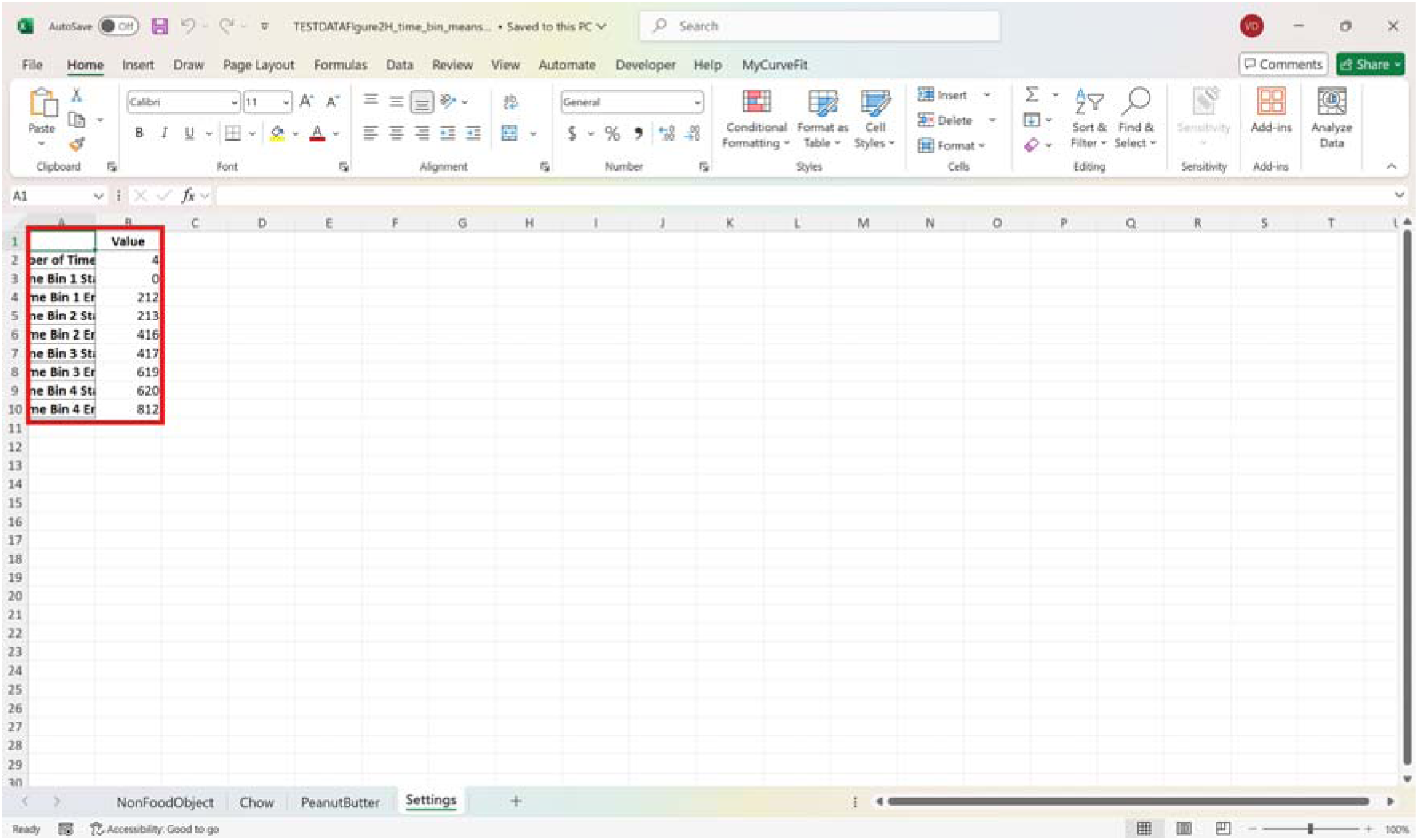

The settings of the number of time bins and the excel row index of each time bin is outputted in the settings sheet of the excel file.

### Bootstrapped confidence intervals and permutation tests

1. Run fiphopha script and select appropriate analysis, in this case it is 2: Within-Groups Bootstrap & Permutation Tests.

2. The following GUI will be presented:

**Box 12.**
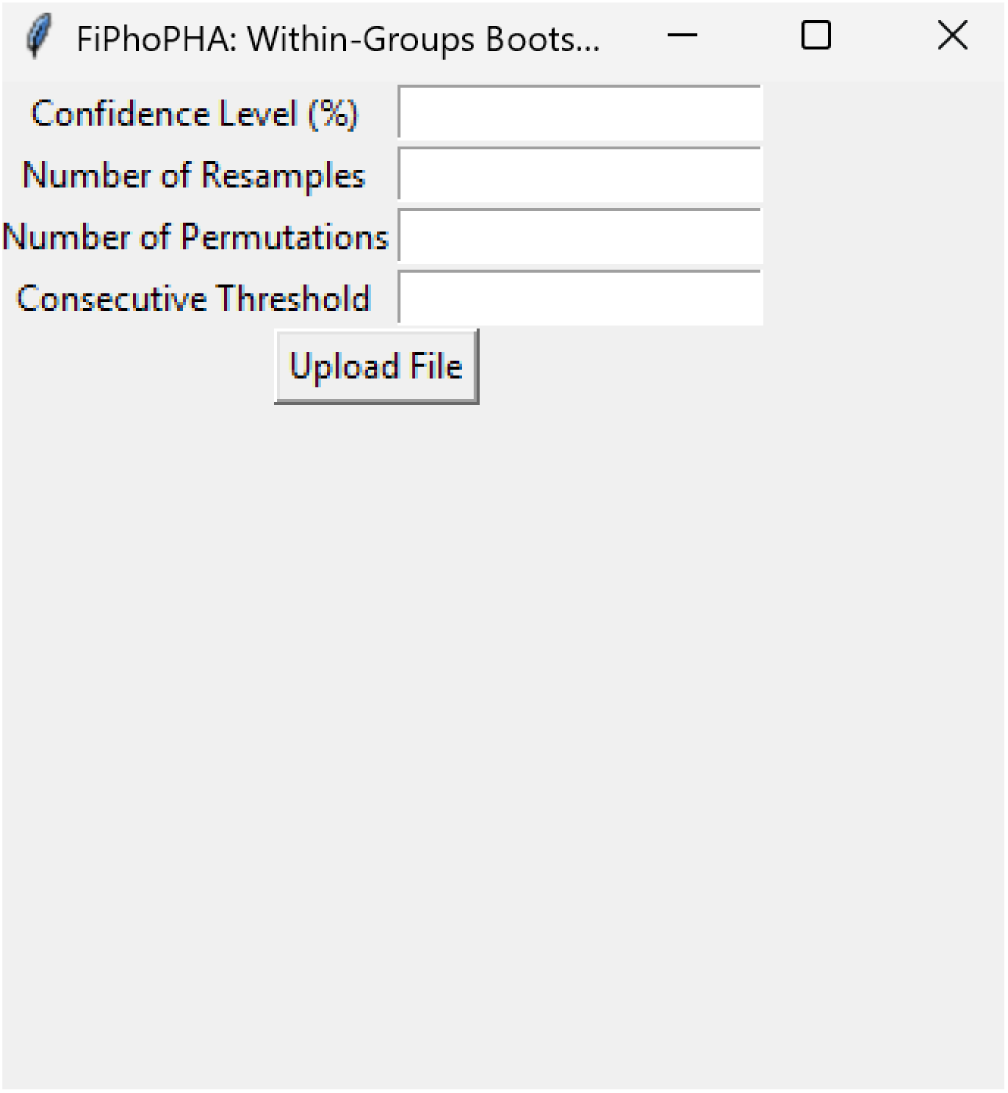

3. Select confidence level, number of resamples, number of permutations, and consecutive threshold, then click upload file and select your input file. We used confidence levels 95% or 99%, and a standard number of 1000 resamples, 1000 permutations based on the previously validated measures of statistical accuracy (Jean-Richard-dit-Bressel et al., 2020). For a consecutive threshold we used 2 (low pass filter), or 4 (kinetics) or 5 (standardised), where our data were sampled at approximately 10 Hz (i.e. every 100 ms), therefore the consecutive threshold represents 2, 4 and 5 sampling periods respectively (i.e. 200, 400 and 500 ms).

4. The following GUI will be shown to select the start row of the baseline, 0 was selected.

**Box 13.**
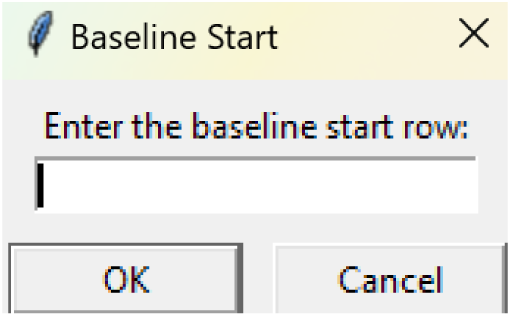

5. Then baseline end is selected, 212 was used.

**Box 14.**
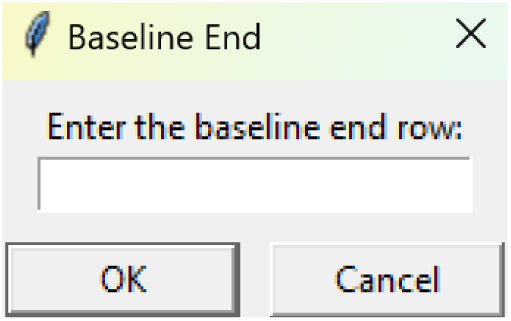

6. After a GUI is shown to enter the start of the comparison period, 213 was used.

**Box 15.**
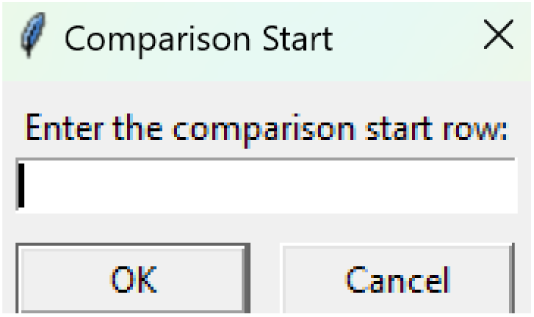

7. A GUI is then shown to enter the end of the comparison period, a max value is shown. This is the maximum row of the data minus 1, as 0 is denoted as row 1 of the excel spreadsheet. 812 was used.

**Box 16.**
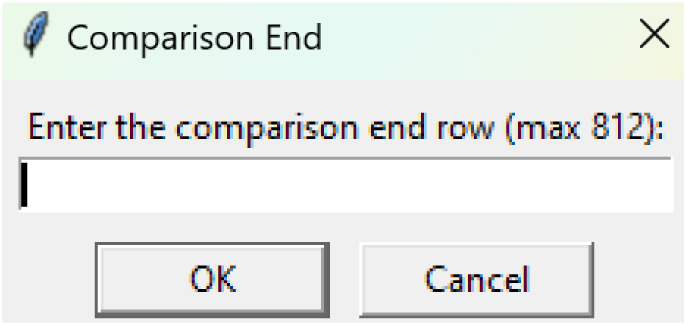

8. An output excel file will be saved to the same directory as the input file with the title of the input file with name_results as shown below.

**Box 17.**
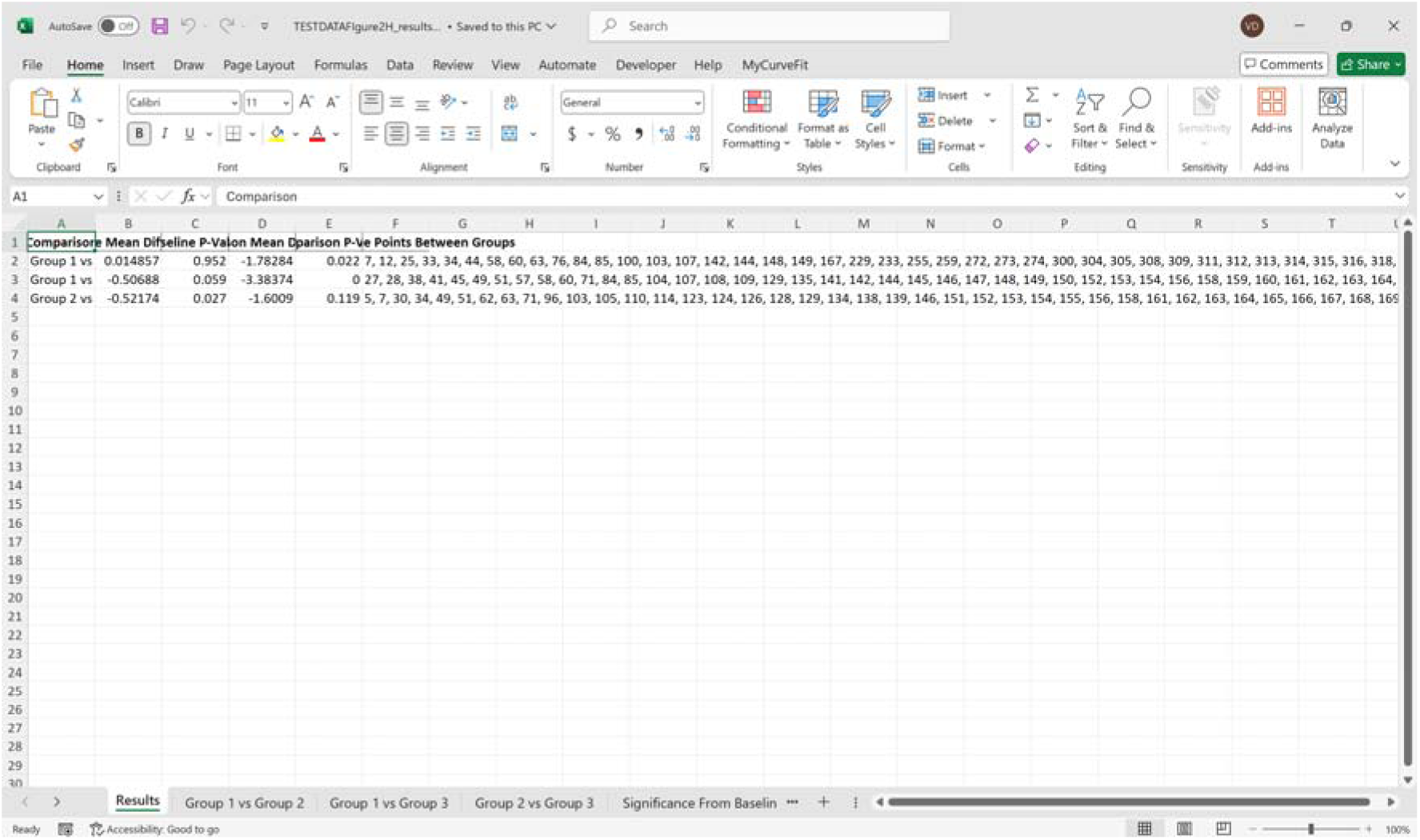

9. This shows the comparison between each group of the input sheet, the mean difference between the groups and p-values of the permutation tests of the defined baseline window, and the same with the comparison period. The last column presents all significant time points between the groups.

**Box 18.**
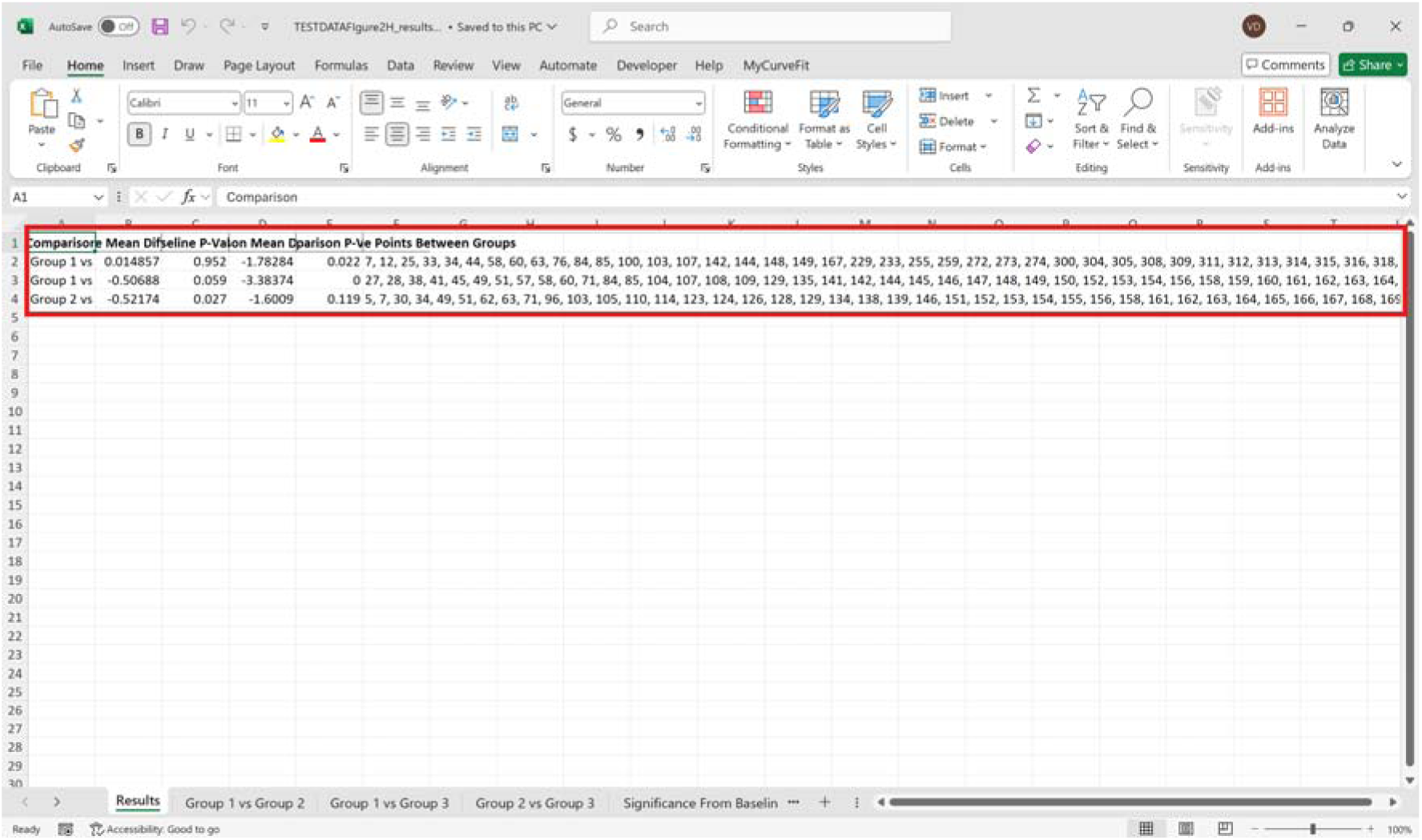

10. This sheet shows a binary significance map of either 0 or 1, with 1 denoting significance for each time point across the group comparisons.

**Box 19.**
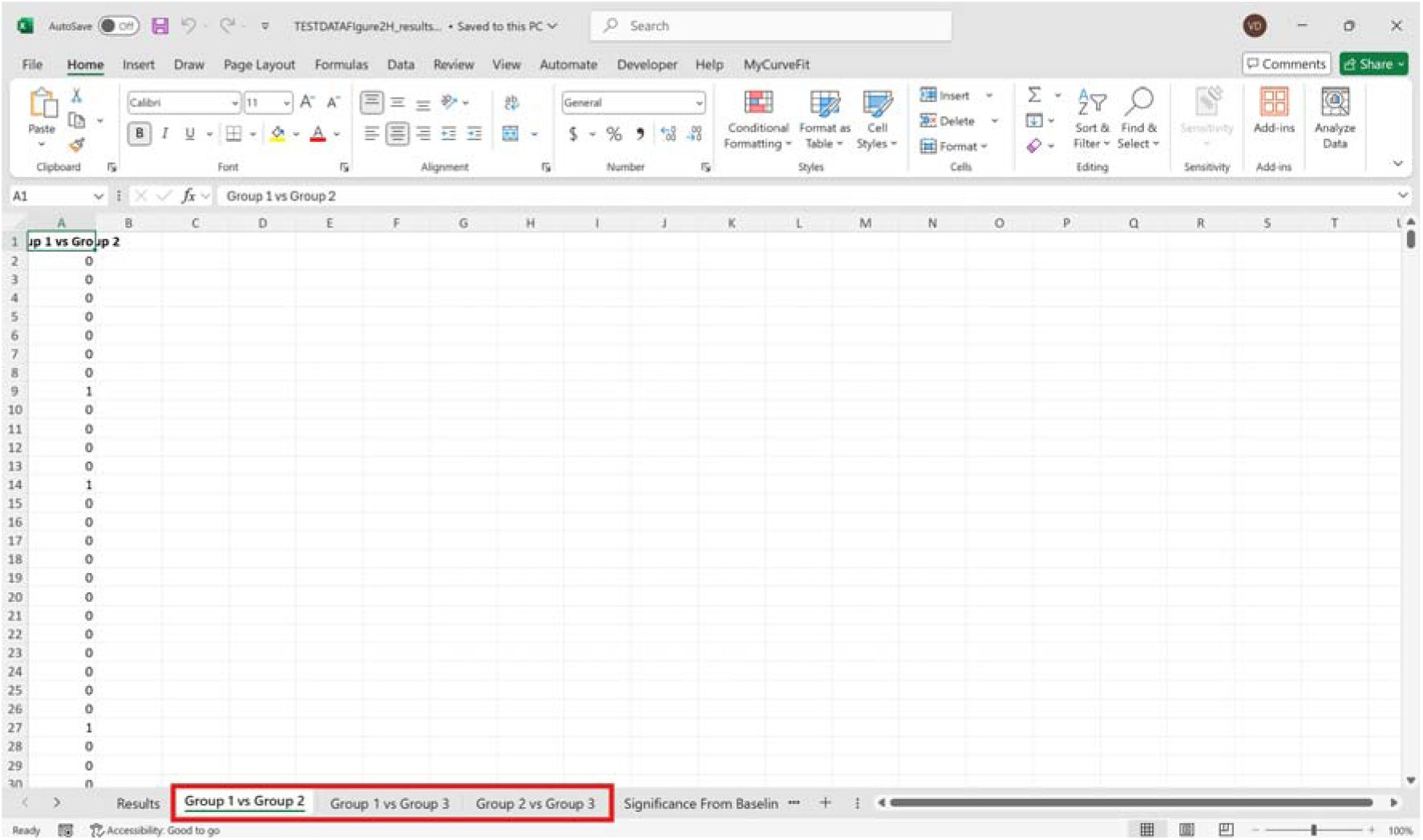

11. This sheet displays the significance from baseline for the groups separately, where 1 denotes significance as shown below.

**Box 20.**
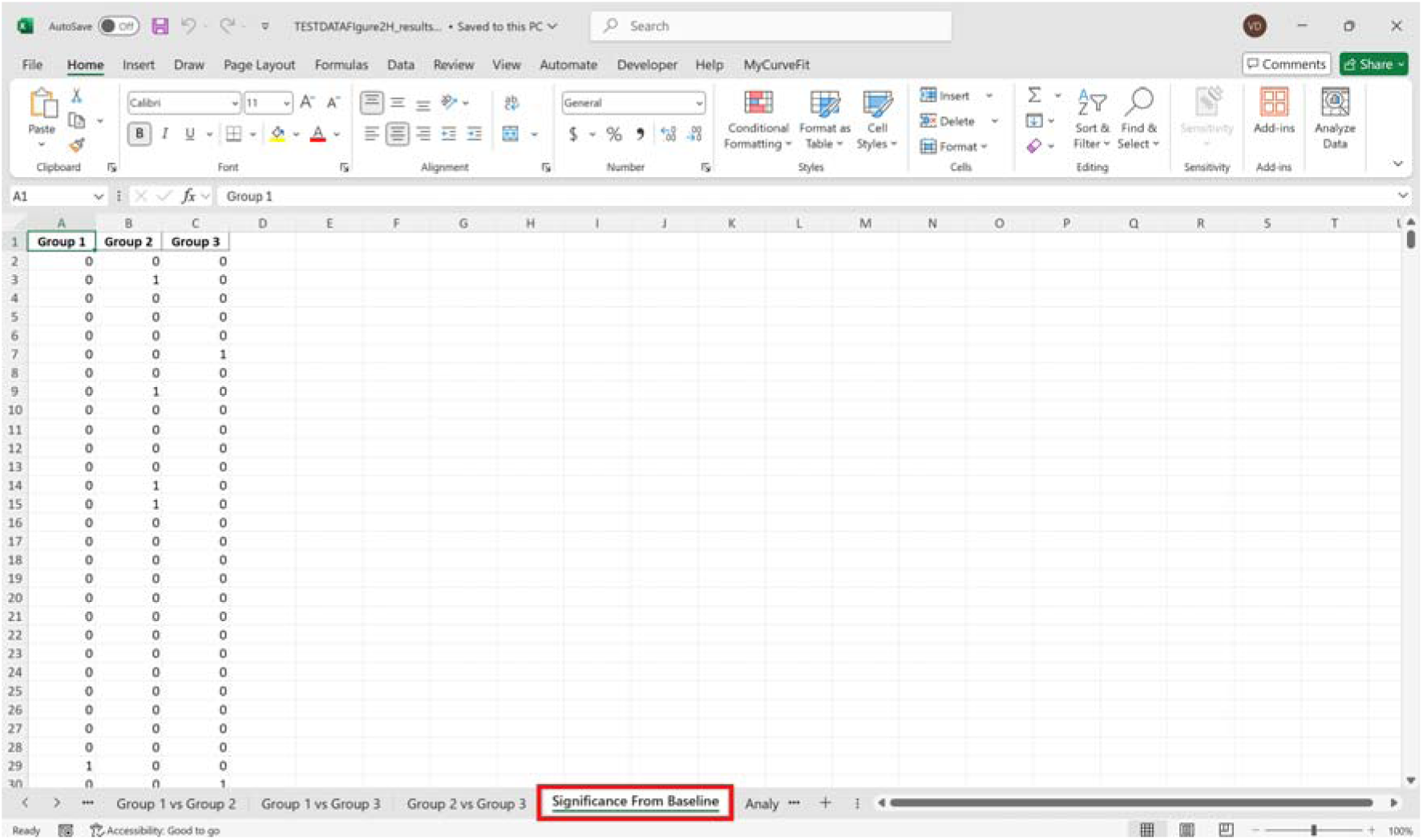

12. The last sheet shows the analysis settings that the user defined in the initial GUI.

**Box 21.**
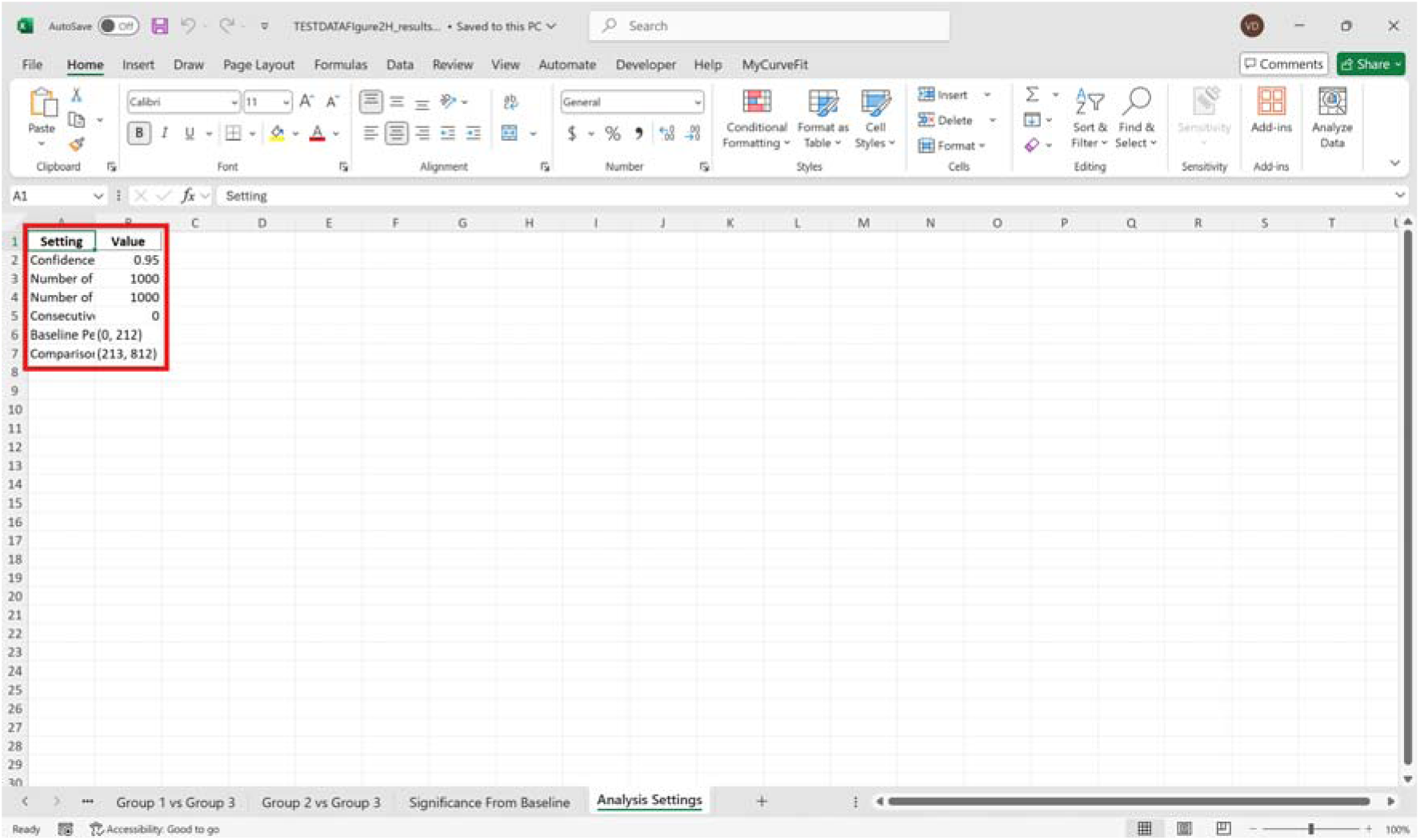

### Common errors

Non-numeric data found in excel sheets or input file is empty.

When picking the row index for analysis, numbers must be non-negative and the data to be analyzed should be equal to or less than the maximum number of rows (refer to box 14-16).

For within-groups analysis, ensure multiple groups have the same number of animals/trials in each excel sheet as each of them are subtracted point by point. If the experiment has been conducted within-groups but due to reasons there are different amounts of animals/trials in each group, it is appropriate to run the between-groups analysis, as it subtracts the bootstrapped distributions of each group.

Input/output directory is linked to a cloud file. The input/output directory must be from hard drive storage on the computer.

## Results

To show the utility of our approach we have re-examined data from Reichenbach and colleagues (Reichenbach et al., 2022), with data representing dopamine release in the nucleus accumbens in overnight fasted wildtype mice with time 0 representing the placement of an object or food into the cage. The data were downsampled to 1Hz, whereas the original data processed at 10Hz (Figure 1A).

**Figure 1.**
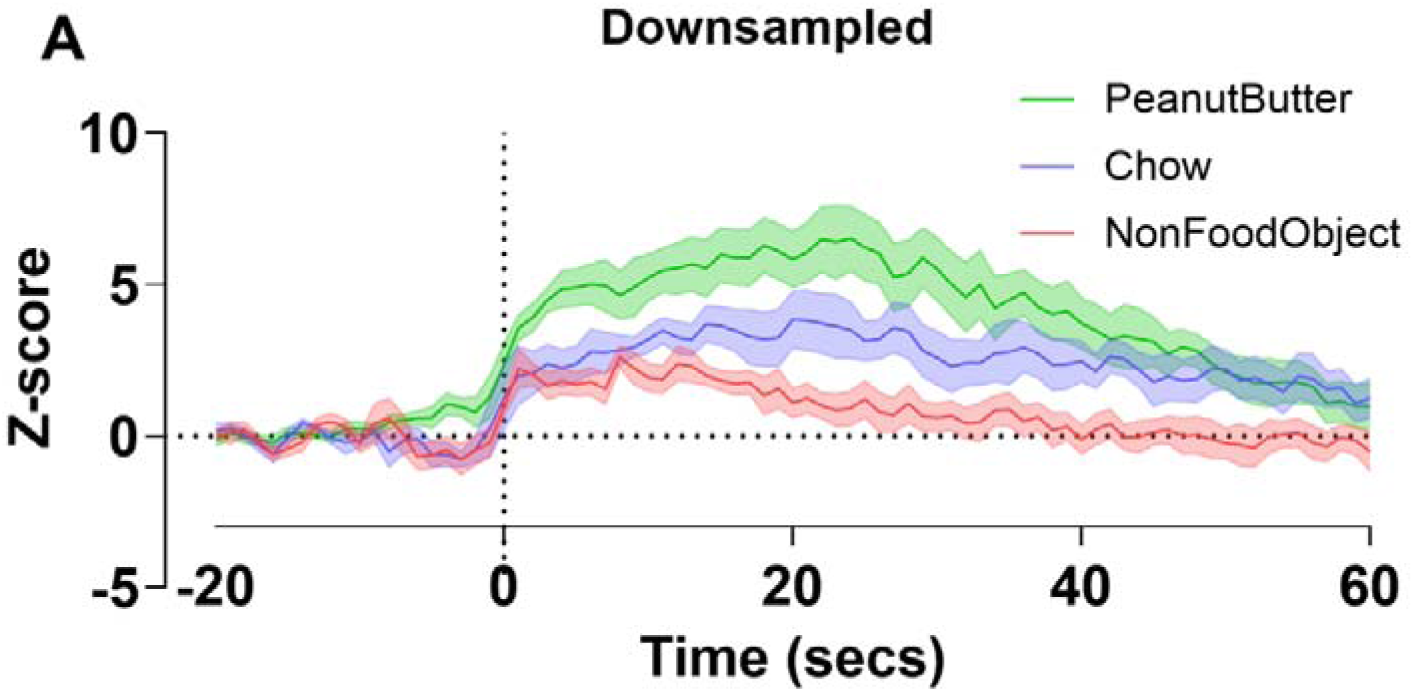
Wildtype overnight fasted nucleus accumbens dopamine release downsampled. Downsampled (by a factor of 10) Z-score photometry trace to approximately 1 Hz for non-food object, chow, and peanut butter put in cage at time 0 (**A**). Data is represented as mean ± standard error of the mean (SEM) and n = 7 in all groups.

### Bootstrapping CIs

#### Differences from baseline

Using the bootstrapped CIs code, we analysed changes in nucleus accumbens dopamine release from baseline after contact with a non-food object, chow, and peanut butter at the original pre-processing of 10Hz (Figure 2A). With 95% CI, individual time points indicate a significance deviation from 0 (baseline) for each group where no consecutive threshold has been applied (Figure 2B1). As the consecutive threshold increases (the amount of timepoints that need to be significant in a row), the amount of timepoints that are deemed significant decreases. At the 95% CI as the threshold increases, no timepoints pre-event are significant at a consecutive threshold of 5 timepoints except for in the peanut butter group (Figure 2B4). The general trends are conserved between the 95% and 99% CI, but at every consecutive threshold there are fewer significant points at 99% CI. At the standardised 5 consecutive points for the 99% CI there are no points in the pre-event baseline period that are significant (Figure 2C4).

**Figure 2.**
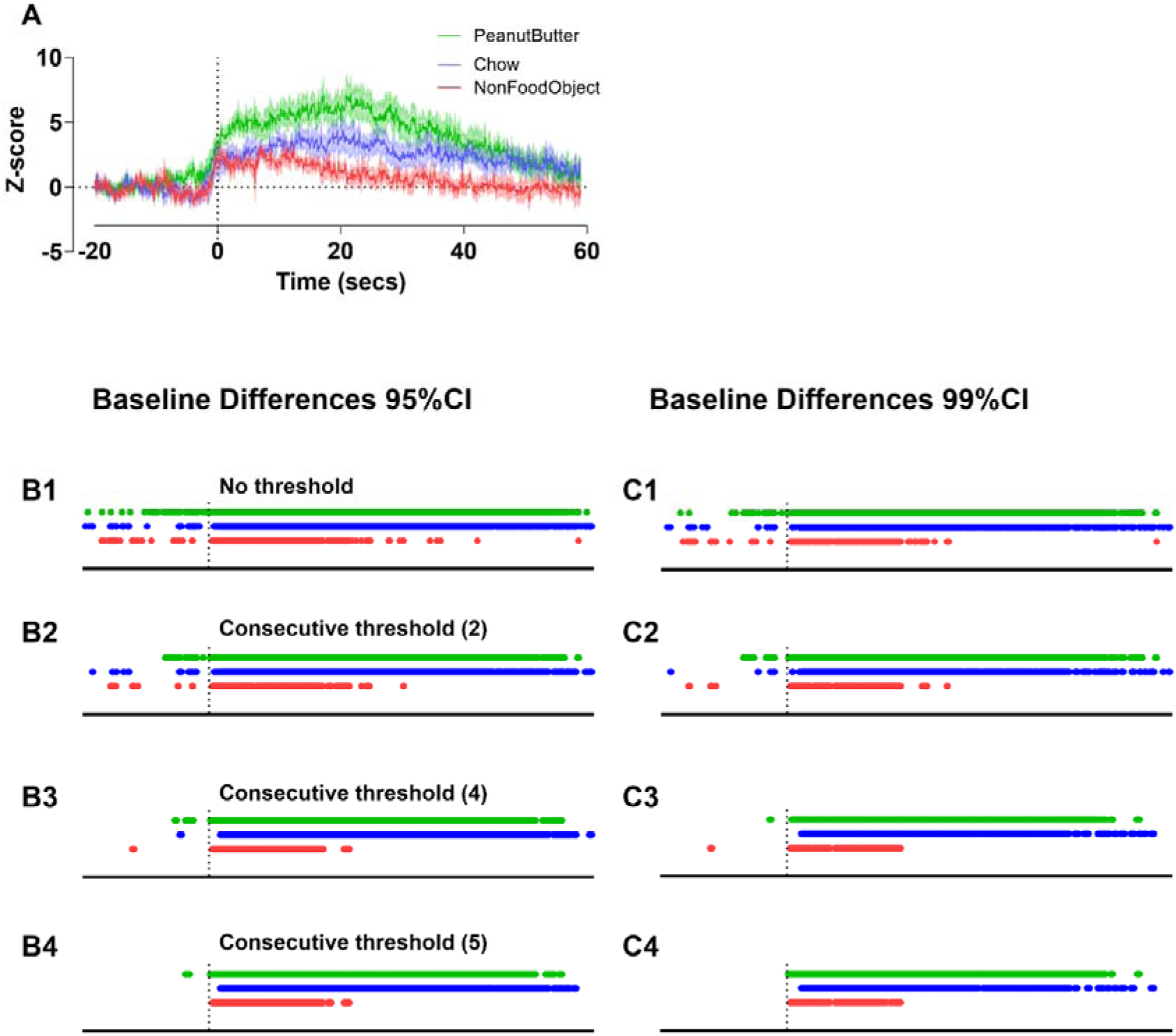
Baseline comparisons in nucleus accumbens dopamine release using bootstrapped CIs with different consecutive thresholds. Original z-score photometry trace for non-food object, chow, and peanut butter put in cage at time 0 (**A**). Baseline differences from bootstrapped CIs with 1000 resamples at 95% confidence with no consecutive threshold (**B1**). Baseline differences from bootstrapped CIs with 1000 resamples at 95% confidence at consecutive threshold 2 (2 significant data points in a row) (**B2**). Baseline differences from bootstrapped CIs with 1000 resamples at 95% confidence at consecutive threshold 4 (4 significant data points in a row) (**B3**). Baseline differences from bootstrapped CIs with 1000 resamples at 95% confidence at consecutive threshold 5 (5 significant data points in a row) (**B4**). Baseline differences from bootstrapped CIs with 1000 resamples at 99% confidence with no consecutive threshold (**C1**). Baseline differences from bootstrapped CIs with 1000 resamples at 99% confidence at consecutive threshold 2 (2 significant data points in a row) (**C2**). Baseline differences from bootstrapped CIs with 1000 resamples at 99% confidence at consecutive threshold 4 (4 significant data points in a row) (**C3**). Baseline differences from bootstrapped CIs with 1000 resamples at 99% confidence at consecutive threshold 5 (5 significant data points in a row) (**C4**). Data is represented as mean ± SEM and n = 7 in all groups. (**A**).

#### Differences between groups

We also analyzed this data using the within-subjects bootstrapped CIs code to assess intergroup comparisons in nucleus accumbens dopamine release (Figure 3A). However, as a point of comparison, we first averaged data into 20 second time bins using the mean time bins code (Figure 3B) and analysed using Two-Way Repeated measured ANOVA with Tukey’s post hoc. This approach is the current standard in the field and gave significant differences between the non-food object and peanut butter during each of the 20 second time intervals after the event and main-effects for object (p<0.01), time (p<0.0001) and an interaction (p<0.001). With 95% CIs with no consecutive threshold (0), all groups showed significant differences during the baseline and post event periods. With no consecutive threshold, the largest intergroup comparisons were in the non-food object vs peanut butter group comparisons (Figure 3C1). This is followed by the non-food object vs chow comparison and then by the chow vs peanut butter comparison. When compared to the results from average time bins (Figure 3B), there is clearly the need for an objective consecutive threshold. The same is shown in the 95% CI at a consecutive threshold of 2, but with fewer timepoints of significance (Figure 3C2). As the consecutive threshold increases to 5, there are fewer points of significance with the same relative trends (Figure 3C4). At the 99% CI, there are overall fewer timepoints of significance for all intergroup comparisons. For this confidence, at a standardised consecutive threshold of 5 timepoints there remains significant differences in all three intergroup comparisons with the non-food object vs peanut butter comparison having the most differences. The other two intergroup comparisons have fewer points of significance that remain less consecutive compared to using lower thresholds (Figure 3D4).

**Figure 3.**
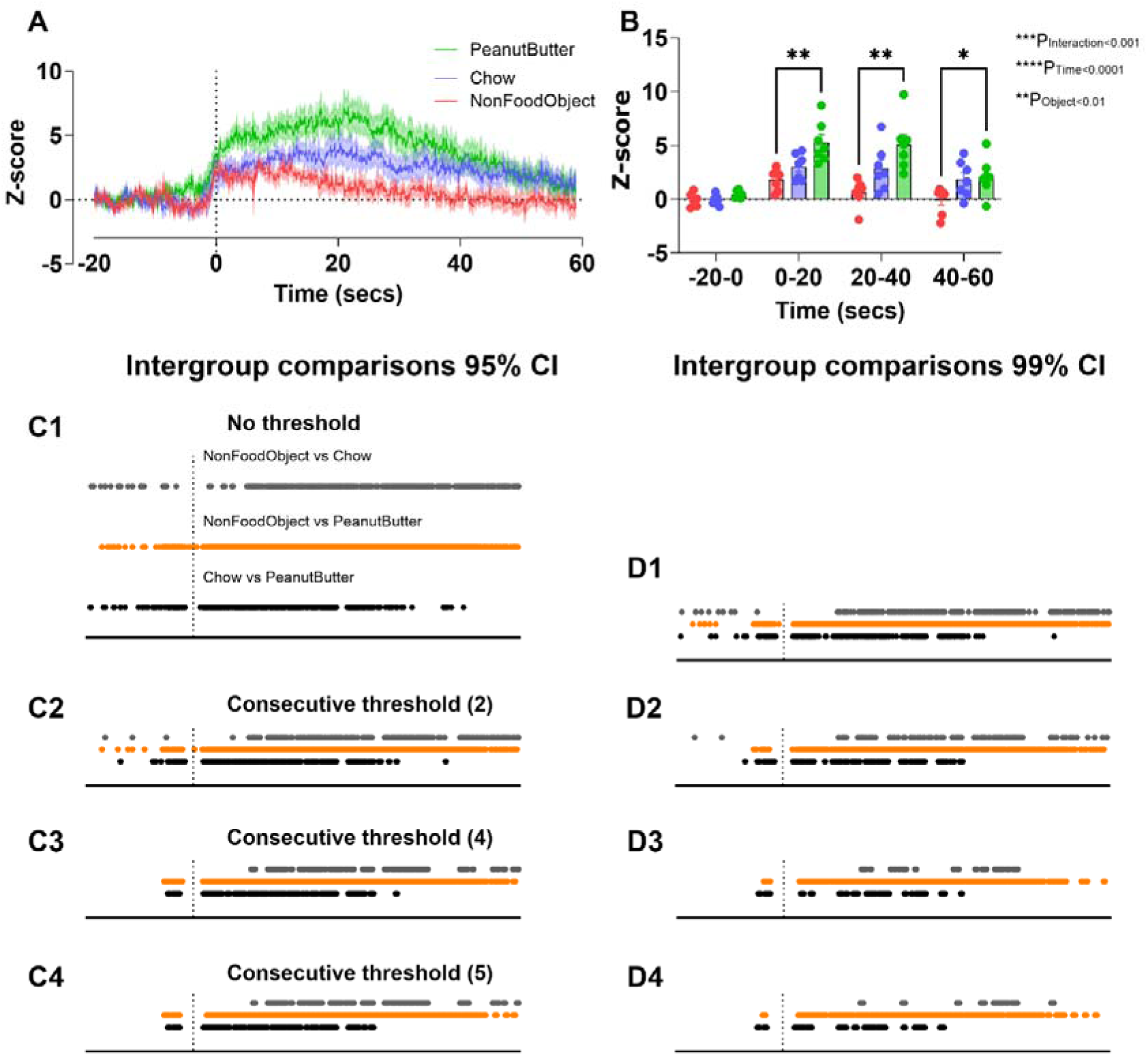
Intergroup differences in nucleus accumbens dopamine release at different consecutive thresholds using bootstrapped CIs. Original z-score photometry trace for non-food object, chow, and peanut butter put in cage at time 0 (**A**). Mean time bins of 20 seconds each of photometry trace with main effects of interaction, time, and object using RM Two-Way ANOVA (**B**). Intergroup differences (non-food object vs chow, non-food object vs peanut butter, and chow vs peanut butter) from bootstrapped CIs with 1000 resamples at 95% confidence with no consecutive threshold (**C1**). Intergroup differences from bootstrapped CIs with 1000 resamples at 95% confidence at consecutive threshold 2 (**C2**). Intergroup differences from bootstrapped CIs with 1000 resamples at 95% confidence at consecutive threshold 4 (**C3**). Intergroup differences from bootstrapped CIs with 1000 resamples at 95% confidence at consecutive threshold 5 (**C4**). Intergroup differences from bootstrapped CIs with 1000 resamples at 99% confidence with no consecutive threshold (**D1**). Intergroup differences from bootstrapped CIs with 1000 resamples at 99% confidence at consecutive threshold 2 (**D2**). Intergroup differences from bootstrapped CIs with 1000 resamples at 99% confidence at consecutive threshold 4 (**D3**). intergroup differences from bootstrapped CIs with 1000 resamples at 99% confidence at consecutive threshold 5 (**D4**). Two-Way Repeated measured ANOVA with Tukey’s multiple comparisons (**B**). Data is represented as mean ± SEM and n = 7 in all groups (**A**) (**B**), *p<0.05, **p<0.01.

### Permutation tests

The permutation tests compare the distributions of the data in either the entire baseline (pre-event) or comparison (post-event) period. Hence changing the consecutive threshold should have little to no effect on these tests. The p-values obtained from the permutation tests represent the probability that the two samples share the same distribution. During the baseline period, p-values change slightly as the consecutive threshold increases (Table 1) at both 95% and 99% CIs.

**Table 1.** Permutation test for baseline (pre-event) differences in nucleus accumbens dopamine release between groups at 95 and 99% confidence intervals.

| Groups | 95% CI-Baseline (Pre-event) p-value |  |  |  |
| --- | --- | --- | --- | --- |
|  | No threshold | Consecutive threshold (2) | Consecutive threshold (4) | Consecutive threshold (5) |
| NonFoodObject vs Chow | 0.952 | 0.948 | 0.951 | 0.957 |
| NonFoodObject vs PeanutButter | 0.059 | 0.061 | 0.058 | 0.068 |
| Chow vs PeanutButter | 0.027 | 0.027 | 0.031 | 0.03 |
|  | 99% CI-Baseline (Pre-event) p-value |  |  |  |
|  | No threshold | Consecutive threshold (2) | Consecutive threshold (4) | Consecutive threshold (5) |
|  | 0.945 | 0.955 | 0.974 | 0.95 |
|  | 0.065 | 0.053 | 0.075 | 0.064 |
|  | 0.041 | 0.027 | 0.027 | 0.032 |

For post-event differences between groups, p-values only change slightly as the consecutive threshold increases (Table 2). The analysis calculates p-values up to three decimal points, hence a p-value of 0, simply means that it is less than 0.001. In general, these results highlight that the various thresholding approaches have little to no effect of permutation test p values. This is because permutations test considers all data points in the baseline or post-event period, where the consecutive threshold approach is not relevant. The slight variation in p-values obtained from the different thresholds is due to the random sampling of data each time. Such that it is unlikely to obtain the same p-values each time you run the permutation tests.

**Table 2.** Permutation test for pre and post-event differences in nucleus accumbens dopamine release between groups using 95 and 99% confidence intervals.

| Groups | 95% CI-Comparison(Post-event) p-value |  |  |  |
| --- | --- | --- | --- | --- |
|  | No threshold | Consecutive threshold (2) | Consecutive threshold (4) | Consecutive threshold (5) |
| NonFoodObject vs Chow | 0.022 | 0.029 | 0.032 | 0.027 |
| NonFoodObject vs PeanutButter | 0 | 0 | 0 | 0.001 |
| Chow vs PeanutButter | 0.119 | 0.119 | 0.104 | 0.104 |
|  | 99% CI-Comparison(Post-event) p-value |  |  |  |
|  | No threshold | Consecutive threshold (2) | Consecutive threshold (4) | Consecutive threshold (5) |
|  | 0.027 | 0.022 | 0.026 | 0.016 |
|  | 0 | 0.001 | 0.001 | 0.001 |
|  | 0.104 | 0.094 | 0.106 | 0.103 |

## Discussion

Currently, statistical analysis of *in vivo* fiber photometry data is unintentionally flawed because it often uses biased user-defined summary statistics for peri-event time bins, such as area under the curve, average z-score, or peak z-score. Therefore, analytical tools to improve statistical reliability in an unbiased manner are essential for data reproducibility. We propose that a standardised easy-to-use post hoc analysis tool with less statistical assumptions would have a beneficial effect on the field. Indeed, with current approaches, changing time bin periods for area under curve analysis of average z-score may change the significance of the results and therefore the conclusions drawn from the experiments. This inadvertently biased approach hinders the field for multiple reasons including different researchers selecting different time bin periods or different parameters to indicate group differences, such as area under the curve, average z-score, max peak z-score. Our bootstrapping CI, based on Jean-Richard-dit-Bressel and colleagues’ approach, removes the need to select the individual parameters or time bin period and thus, provides an unbiased analysis to strengthen the reliability of research outcomes (Jean-Richard-dit-Bressel et al., 2020). Recently, another post-hoc analysis tool was published (Loewinger et al., 2025), however this approach relies on data modelling with important assumptions about data distribution, such as convolving to a linear distribution, which may not be appropriate for the noisy, non-linear, and non-parametric nature of fiber photometry signals. This approach is likely to be computationally demanding, which may hinder its use for large datasets. Regression based approaches have been used previously and can be helpful for modelling fiber photometry data especially when used to study learning across smaller timescales (Mohebi et al., 2019; Lak et al., 2020). Given this approach, it may be useful in conjunction with FiPhoPHA for specific research questions as previously published (Hart et al., 2024). However, perhaps the main advantage of our approach is the usability and ease of adoption that we have intentionally built into the analysis. Our approach does not require a strong coding background and can be simply applied by installing the fiphopha package in python. The graphical user interface enables the user to logically choose and execute the relevant analyses, and the data summary sheets expedite the presentation of statistical differences.

Our reanalysis of data taken from our previous study (Reichenbach et al., 2022) using the bootstrapping confidence intervals highlights greater statistical significance with greater temporal precision than 20 second time bin windows previously analysed. For example, dopamine release to peanut butter is significantly different from the non-food object in both time binned analysis and bootstrapping. However, our new bootstrapping confidence interval analysis highlights significant differences between the non-food object and chow, as well as between chow and peanut butter, which were not observed with our previous time binned analysis. The time bin approach loses the ability to detect significance at precise time periods because time bins condense hundreds of data points into one data point. Our 20 second time bins from Reichenbach et al., (2022) represent condensing 200 data points sampled at 10 Hz into one data point (0.05 Hz) and therefore it is not surprising that our bootstrapping confidence interval analysis has identified additional significance at precise time periods between additional groups. Thus, our new analysis approach has identified that although the conclusions from Reichenbach et al., (2022) are valid, they are not as accurate as they could have been and left important statistical information undiscovered.

For our analysis pipeline, we recommend taking either the entire period before event onset or the entire recording period as the baseline value. Since the bootstrapped CIs have the possibility of type 1 errors, we recommend using the most conservative consecutive threshold measure out of the low-pass filter applied during data collection or the kinetics of the sensor. We have also incorporated analysis for between and within-subjects intergroup comparisons that should be chosen accordingly. Depending on your specific physiological application it may be more appropriate to use a standardised threshold or 99% CI vs 95% CI. Since the 99% CI uses a larger range of confidence of the signal at each individual timepoint, it allows for a more accurate measurement in some cases. In conjunction, the permutation test p-values provide a statistical interrogation of the distribution of data during each period. Furthermore, it is important to not extrapolate beyond the given data, and to recognise that the permutation tests for the baseline and comparison period give an exact p-value, but the number of permutations that is chosen will change the smallest possible p-value. Thus, we suggest that researchers consider increasing the number of permutations when dealing with a smaller sample size. Regardless, coupled with the bootstrapped CIs, this provides measurements of significance from multiple tests.

All analyses were performed on a 13^th^ generation 14 core Intel Core i9-13900H CPU with 16GB of RAM. Most of the tests are not computationally demanding except for the bootstrapped CIs and permutation tests and the extent of the demand is dependent on the amount of data. Multiprocessing is incorporated in the code to utilise all cores of a computer’s processing power, for this reason we recommend running the code with limited background applications/processes concurrently running. The bootstrapped CIs and permutation tests took approximately 1-2mins with the previously described computer specifications, during which, 60-100% of the CPU was used and 70% of RAM. For this case we recommend having a computer with at least 8 GB of RAM. If larger datasets become too computationally demanding, we recommend downsampling the data to a greater extent or reducing the amount of resamples and permutations.

## Conclusion

The various packages available for fiber photometry analysis lack usability, accessibility, and objective post-hoc statistical methods. Here we introduce FiPhoPHA as a python package that performs post-hoc analyses on any raw data set. FiPhoPHA serves as post hoc analysis toolkit to improve reliability and data reproducibility in an unbiased manner for all photometry users. This package incorporates a downsampler, mean time bins, bootstrapped CIs and permutation tests.

